# Parental effects provide an opportunity for coral resilience following major bleaching events

**DOI:** 10.1101/2023.08.09.552721

**Authors:** Elizabeth A. Lenz, Megan J. Donahue, Ruth D. Gates, Hollie M. Putnam, Eveline van der Steeg, Jacqueline L. Padilla-Gamiño

**Author notes:** Deceased (Ruth D. Gates).

## Abstract

Identifying processes that promote coral reef recovery and resilience is crucial as ocean warming becomes more frequent and severe. Sexual reproduction is essential for the replenishment of coral populations and maintenance of genetic diversity; however, the ability for corals to reproduce may be impaired by marine heatwaves that cause coral bleaching. In 2014 and 2015, the Hawaiian Islands experienced coral bleaching with differential bleaching susceptibility in the species *Montipora capitata*, a dominant reef-building coral in the region. We tested the hypothesis that coral bleaching resistance enhances reproductive capacity and offspring performance by examining the reproductive biology of colonies that bleached and recovered (B) and colonies that did not bleach (NB) in 2015 in the subsequent spawning seasons. The proportion of colonies that spawned was higher in 2016 than in 2017. Regardless of parental bleaching history, we found eggs with higher abnormality and bundles with fewer eggs in 2016 than 2017. While reproductive output was similar between B and NB colonies in 2016, survivorship of offspring that year were significantly influenced by the parental bleaching history (egg donor × sperm donor: B × B, B × NB, NB × B, and NB × NB). Offspring produced by NB egg donors had the highest survivorship, while offspring from previously bleached colonies had the lowest survivorship, highlighting the negative effects of bleaching on parental investment and offspring performance. While sexual reproduction continues in *M. capitata* post-bleaching, gametes are differentially impacted by recovery time following a bleaching event and by parental bleaching resistance. Our results demonstrate the importance of identifying bleaching resistant individuals during and after heating events. This study further highlights the significance of maternal effects through potential egg provisioning for offspring survivorship and provides a baseline for human-assisted intervention (i.e., selective breeding) to mitigate the effects of climate change on coral reefs.

## INTRODUCTION

Ocean warming caused by anthropogenic greenhouse gas emissions is one of the primary threats to the function of shallow tropical coral reefs (Gattuso et al., 2015; Intergovernmental Panel on Climate Change, 2018). Prolonged warming above the local thermal threshold for bleaching coupled with high irradiances can cause severe coral bleaching (Glynn, 1996), the disruption of the nutritional symbiosis between the coral host and its unicellular dinoflagellates, Symbiodiniaceae (formerly, *Symbiodinium* spp.; LaJeunesse et al., 2018). This can subsequently result in increased rates of disease transmission (Muller et al., 2018) and mortality (Hughes et al., 2018b) along with reduced calcification rates and reproductive capacity in corals (Szmant & Gassman, 1990; Baird & Marshall, 2002). Continual declines in coral cover are predicted given the range of local and global disturbances simulateneously acting on coral reefs, with warming ranked as the most severe (Gardner et al., 2003; De’ath et al., 2012; Hughes et al., 2018a). Identifying sources of resilience in coral reef ecosystems, such as locating exceptional coral genotypes that can thrive under extreme warming or temperature fluctuations, will be key in maintaining and restoring reefs for the future.

Differential bleaching susceptibility (Cunning et al., 2016; Loya et al., 2001; Wall et al., 2019) during a thermal stress event illustrates biological variation within populations that may serve as a source of resilience and an opportunity for selection through reproductive success (Barott et al., 2021; Johnston et al., 2020). Thermal tolerance and capacity to recover after bleaching are important factors that influence sexual reproduction, recruitment, and success of future generations to adapt (Szmant & Gassman, 1990; Baird & Marshall, 2002; Levitan et al., 2014; Putnam, 2021). Successful sexual reproduction and recruitment are essential in maintaining coral populations (Bramanti & Edmunds, 2016), repopulating disturbed coral reefs (Bellwood et al., 2004; Gilmour et al., 2013; Adjeroud et al., 2017; Cruz & Harrison, 2017), and enhancing genetic diversity within populations to overcome selective pressures (Richmond, 1997; van Oppen & Gates, 2006). However, parental investment in gametogenesis is energetically costly (Vance, 1973), and for corals reproductive cycles may exceed six to ten months (Richmond & Hunter, 1990; Padilla-Gamiño et al., 2014). Therefore, prolonged environmental stress can drive prioritization of energetic investment into basic metabolic function and repair, at the expense of growth and sexual reproduction (Richmond, 1987; Ward, 1995; Leuzinger et al., 2012). Importantly, this tradeoff in energetic investment is likely to depend on the susceptibility and severity of coral bleaching, with greater energy available for reproduction in corals resistant to bleaching (Lenz et al., 2021).

Coral bleaching is known to impact sexual reproduction (Baird & Marshall, 2002; Fisch et al., 2019) and recruitment (Hughes et al. 2019; Price et al., 2019). For example, after the 1987 coral bleaching event in the Caribbean, *Orbicella annularis* recovered from bleaching by metabolizing tissue biomass, but did not complete gametogenesis in the following months, whereas colonies that had not bleached of the same species were able to develop and release gametes (Szmant & Gassman, 1990). Similarly, during the 1998 bleaching event on the Great Barrier Reef, bleached corals showed high variation in reproduction compared to colonies resistant to bleaching nearby that experienced the same thermal stress. For acroporid species, reproductive polyps were more common in colonies that did not bleach, with larger eggs at higher densities per polyp than colonies that bleached and recovered (Ward et al., 2002).

Given logistical complexities and challenges, most studies have primarily investigated gametogenesis in the life cycle of coral with some understanding of cross-generational effects (i.e., parental, carry-over, or transgenerational effects) following major bleaching events. The impacts of coral bleaching may last for months to years after the initial thermal stress (Hagedorn et al., 2016), and can manifest in life stages downstream such as fertilization (Negri et al., 2007; Howells et al., 2016; Omori et al., 2001), larval development, and recruitment (Edmunds, 2018; Hughes et al. 2019; Price et al., 2019). Between the 2005 and 2010 bleaching events in Panama, Levitan et al. (2014) found that thermally tolerant *Orbicella franksi* recovered the capacity to produce and release gametes more quickly (within 3 to 5 years) than the more thermally sensitive *O. annularis*. In Mo’orea, French Polynesia following a minor coral bleaching in 2016 during El Niño, there was high coral recruitment indicating that there was a source population that maintained reproductive function (Edmunds 2017). While these studies demonstrate a range of responses in sexual reproductive biology and ecology during recovery post bleaching (i.e., gametogenesis and recruitment), few studies have followed both the intra- and intergenerational impacts of bleaching. Recent marine heatwaves eliciting differential coral bleaching of *M. capitata* in Hawaiʻi provide an opportunity to compare the impacts of parental bleaching history on coral reproduction and offspring performance during recovery and offer potential insight on coral resilience (Barott et al., 2021; Dilworth et al., 2021; Drury et al., 2022).

Coral reefs in the subtropical waters of Hawaiʻi were largely naive to global bleaching events (Jokiel & Coles, 1990; Jokiel & Brown, 2004; Bahr et al., 2016) with bleaching events first recorded in the Main Hawaiian Islands in 1996 and then in the Northwestern Hawaiian Islands in 2002 (Jokiel & Coles, 1990; Jokiel & Brown, 2004; Bahr et al., 2016). However, the Hawaiian Archipelago experienced “the blob” heatwave, followed by an El Niño that resulted in severe back-to-back coral bleaching in 2014 and 2015 (Figure 1A; Bahr et al., 2017; Couch et al., 2017) (Figure 3). During these consecutive bleaching events, degree heating weeks (DHW) in the Main Hawaiian Islands exceeded 8 weeks by September in both years (Bahr et al., 2017; Couch et al., 2017). In Kāneʻohe Bay (Oʻahu, Hawaiʻi), ∼70% of reef corals on the shallow reefs (< 2 m depth) bleached and exhibited 13 - 22% mortality in 2014 and 2015 (Bahr et al., 2015; Bahr et al., 2018; Coles et al., 2018; Rodgers *et al*., 2017). During both events in Kāneʻohe Bay, colonies of the dominant reef-building coral, *Montipora capitata*, visibly bleached or remained pigmented during prolonged heat stress (Figure 1B). Despite widespread bleaching, approximately 70% of *M. capitata* that bleached in 2014 and 2015 were considered recovered by the following December and January based on visual coloration (Bahr et al, 2018; Cunning et al., 2016; Wall et al. 2019; Matsuda et al., 2020; Barott et al., 2021).

**Figure 1.**
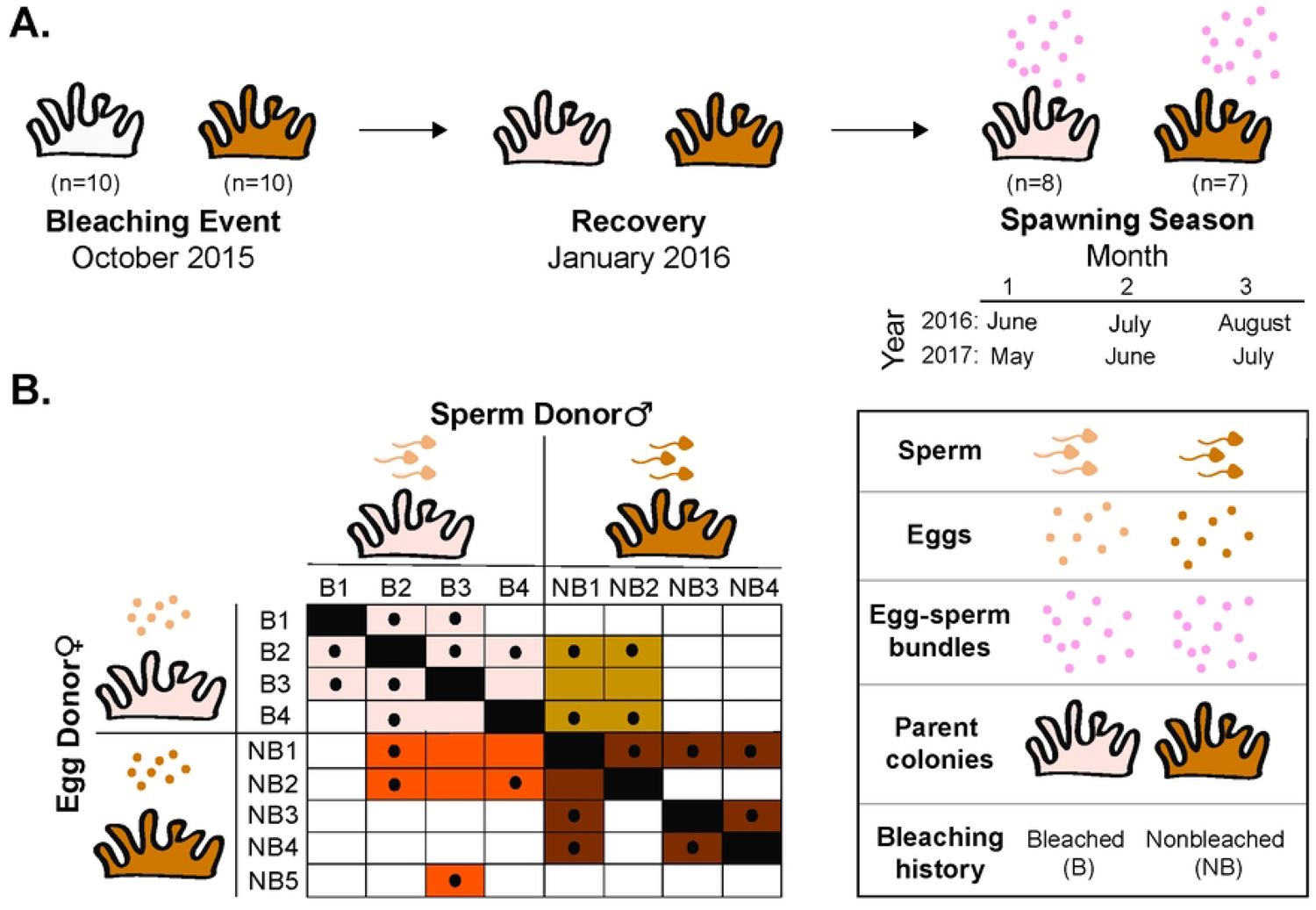
A) Temperature data from 2010 to 2017 (NOAA Buoy Moku oʻ Loe, HI Station ID: 1612480) illustrate historical patterns and identify years of bleaching events in Oʻahu, Hawaiʻi. The bleaching threshold between 30 to 31 °C of corals in Kāneʻohe Bay (Coles et al., 2018) is shown in the shaded red, thermometers indicate the 2014 and 2015 bleaching events and the spawning corals indicate the spawning seasons. B) An image depicting the tagged bleached (left) and nonbleached (right) parental colonies in response to the 2015 heat stress in Kāneʻohe Bay.

*M. capitata* demonstrates relatively high tolerance against multiple local and global stressors (Gibbin et al., 2015; Putnam et al., 2016), with varied sensitivity among individual colonies and their traits measured under elevated temperature (Matsuda et al., 2020; Barott et al., 2021), such as survivorship (Coles et al., 2018), growth (Bahr et al., 2016), and biomass composition (Wall et al. 2019; Grottoli et al., 2004, 2006; Rodrigues & Grottoli, 2007; Bahr et al., 2016). Reproductive effort of *M. capitata*, particularly oocyte characteristics and spawning, has shown little response to warming (Cox, 2007; Hagedorn et al., 2016). This reproductive response may contribute to its ecological success along the fringing and patch reefs of Kāneʻohe Bay (Kolinski, 2004). However, percent of motile sperm from *M. capitata* declined from 80-90% in 2011 to 40.5% in 2015, corresponding with the consecutive bleaching events in Kāneʻohe Bay (Hagedorn et al., 2016). For *M. capitata,* oogenesis can begin as early as July, which means that early egg development may cooccur with severe, prolonged warming events (July-October), and later egg development continues when corals are recovering from these events (Nov-June to August). This could create a strain on energetic resources when corals are compromised during a substantial fraction of the typical gametogenic cycle (Padilla-Gamiño & Gates, 2012; Padilla-Gamiño et al., 2014). Therefore, tracking *M. capitata* through subsequent spawning seasons after bleaching events can reveal the reproductive capacity of this species as ocean temperature continues to increase.

In this study, we examined cross-generation plasticity (i.e., parental effects) to determine how parental response to environmental events influence reproduction (Byrne et al., 2019). We measured the reproductive biology of *M. capitata* for two spawning seasons (2016 and 2017) following bleaching events (2014 and 2015). We tested the following hypotheses: (i) that parental bleaching history [bleached (B) and nonbleached (NB)] would affect reproductive performance in subsequent spawning seasons and (ii) intentional crosses of gametes from parent colonies of differential bleaching history would influence offspring success (Figure 2A). In 2016, we tested the second hypothesis and quantified the downstream effects of parental bleaching history from gamete release to settlement of the offspring in parent colonies that did and did not bleach during the 2015 warming event (Figure 2B). This study was designed to assess selective processes in nature confronted by climate change while also exploring breeding techniques as an intervention strategy for coral restoration to maintain genetic diversity.

**Figure 2.**
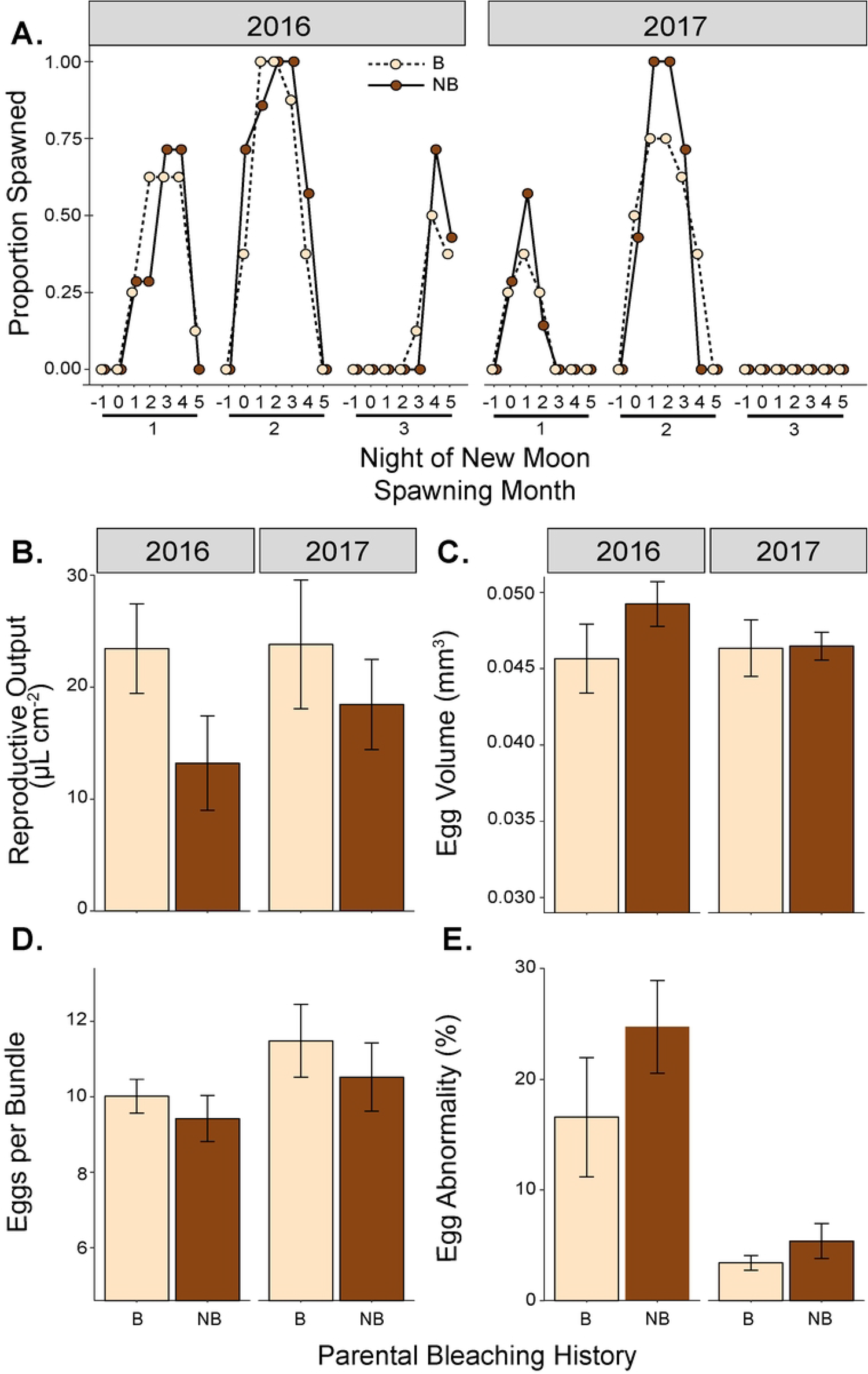
Experimental design of the study. A) Bleached and nonbleached colonies were tagged in October 2015 at the peak of the bleaching event. Bleached colonies in this experiment recovered by January 2016. Total reproductive output and gamete collections were measured during the 2016 and 2017 spawning seasons. Months of the spawning season differ between years because of the different timing of the new moon in 2016 and 2017. B) Selective breeding matrix illustrating the crossing of egg and sperm donors conducted in July 2016 based on parental bleaching history. Colored squares indicate the cross of individuals attempted and solid black circles indicate successful fertilization. Offspring from these crosses were used to measure survivorship of larvae and settlers and settlement.

## MATERIALS AND METHODS

### 2.1 Selecting parent colonies and spawning events

*Montipora capitata* is a hermaphroditic broadcast spawner and its reproductive cycle, spawning dynamics, and early life stages have been extensively studied at the Hawai‘i Institute of Marine Biology (HIMB) located in Kāneʻohe Bay, on the windward side of Oʻahu, Hawaiʻi, USA (Richmond & Hunter, 1990; Padilla-Gamiño et al., 2011, 2012; Padilla-Gamiño & Gates, 2012; Hagedorn et al., 2015). In Hawaiʻi, oogenesis begins a 9–10 month period as early as July and as late as October, while spermatogenesis begins the following April to May, ca. 1 month prior to the first spawning event in May or June (Padilla-Gamiño et al., 2014), creating the potential for differential effects of bleaching on oocytes and sperm. Symbiodiniaceae are vertically transferred from *M. capitata* parent colonies into eggs prior to the formation of the egg-sperm bundles, which are released during spawning (Padilla-Gamiño et al., 2011). Spawning in *M. capitata* extends over three, consecutive lunar months between May and September for 3 to 5 consecutive nights between 20:45 and 22:30 hrs, starting on the night of the new moon (Richmond & Hunter, 1990; Padilla-Gamiño & Gates, 2012). The second and third nights are when the largest spawning events most commonly occur (Padilla-Gamiño & Gates, 2012).

During the peak of the 2015 bleaching event in Hawaiʻi, ten pairs of colonies (30-100 cm diameter) of *M. capitata* were identified and tagged as bleached (B) and nonbleached (NB) along the leeward side of the reef surrounding HIMB (21’ 26.09 N, 157’ 47.47’ W) on 20 October 2015 (Figure 3C). These colonies remained in the field until retrieved three days prior to the new moon of the spawning months in 2016 (June, July, and August) and 2017 (May, June, and July) (Figure 3A). To examine reproductive performance of B and NB colonies of *M. capitat*a, parent colonies were collected by removing the entire colony from the reef, or by breaking large fragments (30-40 cm in diameter) from tagged colonies using a hammer and chisel. These collections were first completed on 4 and 5 June 2016. Of the twenty colonies tagged, seven colonies that had not bleached and eight colonies that had bleached and recovered were alive and used for the study. The other five colonies not recovered had either died or were missing from the reef. The fifteen colonies were transported to the wet laboratory at HIMB in 20-L buckets filled with seawater from Kāneʻohe Bay at an ambient temperature of ∼28 to 29 °C. Colonies were randomly allocated to two ∼1,300-L shaded outdoor flow-through tanks (Putnam et al., 2016; Gibbin et al., 2018). Both tanks had sand-filtered seawater delivered at a flow rate of ∼6-L minute^-1^ and a circulation pump (700 gph Magnetic Drive, Danner Manufacturing Inc. Islandia, NY, USA). Irradiance and temperature within each tank were recorded every fifteen minutes with a cosine corrected photosynthetically active radiation (PAR) sensor (Odyssey PAR loggers, Dataflow Systems Ltd, Christchurch, NZ) calibrated to a Licor 192SA sensor, and a temperature logger (Hobo™ Water Temp Pro v2 resolution ± 0.2°C, Onset Computer Corporation, Bourne, MA, USA). Three to five days after each spawning event, colonies were returned to the original field site by attaching them to a fixed rack with cable ties and retrieved two days before the next new moon of the spawning season.

**Figure 3.**
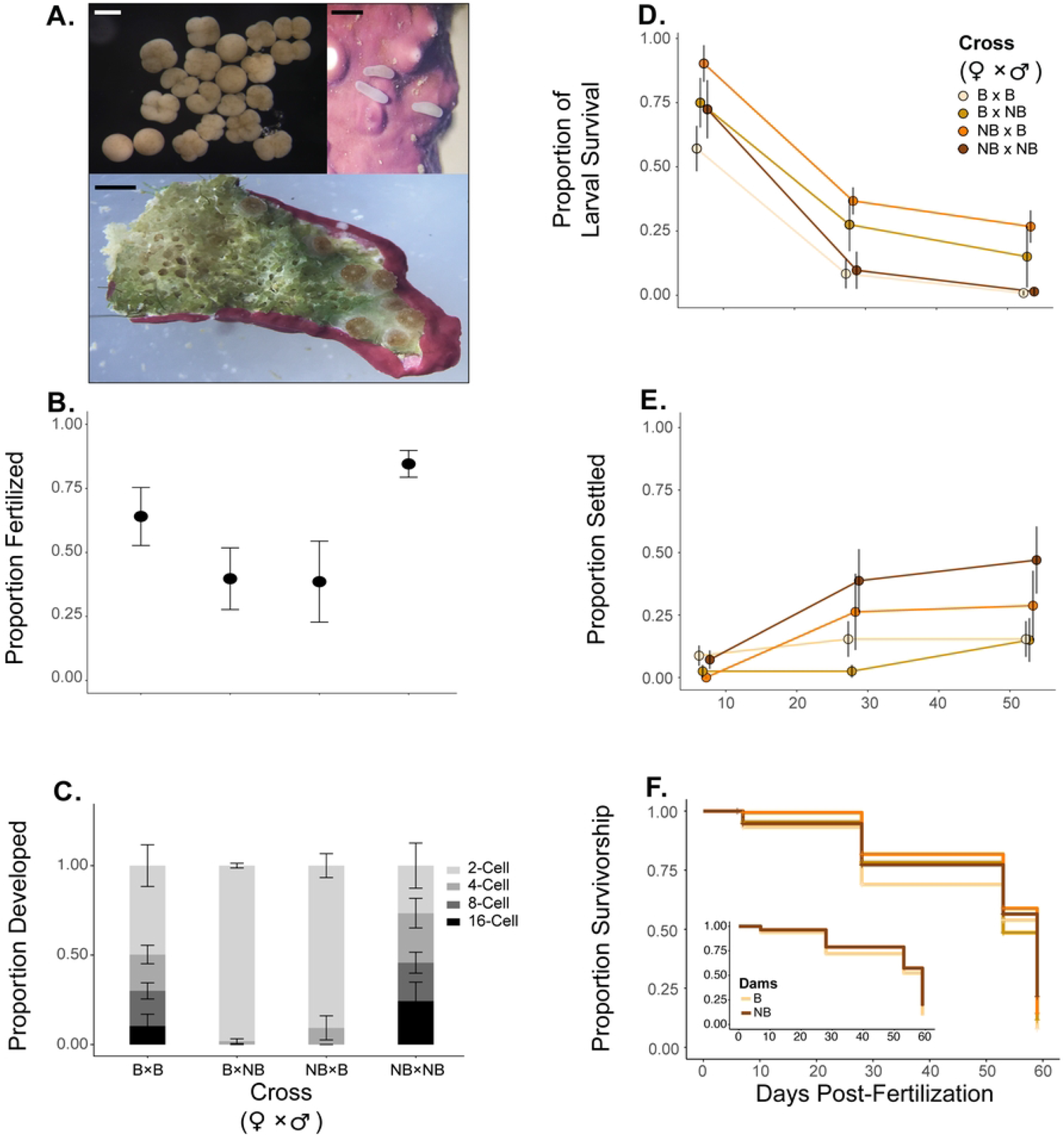
Reproductive traits measured from the same parent colonies in Fig. 2 during 2016 and 2017 spawning seasons following the 2015 bleaching event. A) Proportion of spawning each night in 2016 and 2017 spawning seasons from parent colonies that had bleached and not bleached. “0” indicates the night of the new moon. Mean (± SE) values for B) reproductive output normalized to planar surface area, C) egg volume, D) number of eggs per bundle, and E) percent egg abnormality measured from bleached and nonbleached parents in 2016 and 2017.

### 2.2 Sexual Reproduction

Starting one night prior to the new moon, *M. capitata* parent colonies were monitored for seven nights. During each night of spawning, colonies were isolated at 19:30 in individual containers filled with ambient seawater from the flow-through tanks. When spawning occurred, *M. capitata* released egg-sperm bundles into the water column between 20:45 and 22:30 with peak spawning typically expected on the second night of the new moon (Richmond & Hunter, 1990; Cox, 2007; Padilla-Gamiño & Gates, 2012). Spawning activity of individual colonies was monitored each night and recorded as “spawn” or “no spawn”. For the spawning colonies, we quantified the total volume of gametes released, number of eggs per bundle, and egg quality (i.e., area and abnormality).

Sterilized disposable pipets (2 mL) were used to gently collect all egg-sperm bundles at the water surface from each individual colony to avoid cross contamination or prematurely breaking the egg-sperm bundles. We preserved 3-5 egg-sperm bundles per colony per night to quantify the number of eggs per bundle, egg volume for size, and abnormality. Each egg-sperm bundle was placed in a 2 mL microcentrifuge tube and allowed to break up in 0.1 mL of seawater and for the eggs to hydrate for 2 hrs before preserved in zinc fixative (1:4 Z-fix, Sigma-Aldrich Inc. to 0.2 µm filtered seawater FSW). Preserved eggs from each bundle were photographed using an Olympus SZX7 dissecting microscope equipped with an Olympus America camera (SN: BH039933-H); from photographs, we counted the number of eggs per bundle and measured the egg diameter using ImageJ2 software (Schneider et al., 2012). Egg volume was calculated using the equation for a sphere with the measured egg diameter of spherical eggs. We also recorded the proportion of abnormal (irregular) eggs packaged within each bundle (Padilla-Gamiño et al. 2011; Hagedorn et al., 2016). Remaining egg-sperm bundles from each colony were placed into individual 50 mL Falcon tubes to quantify the total volume of gametes of each colony per night. Annual reproductive output per colony was estimated by summing the spawn volume across the entire spawning season, normalized to planar surface area of the colony using Fiji software (Schindelin et al., 2012).

### 2.3 Fertilization success and larval survivorship

To compare offspring performance of bleached and nonbleached parents, we isolated the egg-sperm bundles from each parental colony that released more than 1 mL of spawn volume on the nights of 5 and 6 July 2016 (peak spawning) and placed egg-sperm bundles from each colony into a separate 50 mL falcon tube. Within one hour of the bundle breaking apart, eggs floated to the surface and sperm sank to the bottom. Sperm were pipetted from the bottom of the tube, and eggs were rinsed twice with 0.2 µm filtered seawater (FSW). Sperm from each colony was placed in separate 50 mL falcon tubes and later used to fertilize eggs from specific colonies.

Nine colonies had adequate spawn volume to include in crosses, and thirty individual crosses were made from gametes based on parental bleaching history to generate four cross-types (egg donor × sperm donor): B × B (n=8), B × NB (n=4), NB × B (n=4), and NB × NB (n=7) (Figure 2B). For fertilization, the eggs (1 mL) were in a concentration of ∼10^6^ sperm mL^-1^ (by visual inspection) within a 50-mL falcon tube (Willis et al., 1997). Thirty minutes after sperm and eggs were mixed, each cross type of fertilized eggs was transferred into individual 1 L conical tanks filled with UV-sterilized 1-µm FSW to avoid polyspermy. For *M. capitat*a, self-fertilization is extremely rare (Hodgson, 1988; Maté et al., 1997). To estimate fertilization success, three subsamples of 20-30 eggs were collected from each conical after approximately 3-hrs (i.e., when initial cleave stages are expected [Babcock & Heyward, 1986; Willis et al., 1997]), placed in a 2-µL microcentrifuge tube, and preserved in Z-fix (1:4 Z-fix to FSW). Remaining embryos in the conical tanks developed, and slow flow rate of FSW was introduced to mitigate potential effects of montiporic acid (Hagedorn et al., 2015). Five days post-fertilization, 10-15 larvae per conical tank were placed in a 10-mL well-plate filled with 5-mL of FSW with a chip of crustose coralline algae to track settlement through time; FSW was exchanged every other day. The proportion of planulae and settlers were examined on days 7, 28, and 53 post-fertilization while the total number of offspring alive were counted on days 6, 7, 28, 53, and 59 post-fertilization to estimate survivorship probability curves.

### 2.4 Statistical Analysis

All analyses were conducted in R (R Core Team, 2014; v. 3.5.1). We used a generalized linear mixed effects model to determine the effects of bleaching history on spawning activity of the 8 B and 7 NB parental colonies observed, we used a generalized linear mixed effect model (*glmer* in *lme4*: Bates et al. 2015) with a binomial response (spawn/no spawn). Bleaching history (B/NB) and year (2016/2017) were included as fixed effects, and spawning month (1/2/3) and colony ID were included as random effects. To analyze total reproductive output, egg size, number of eggs per bundle, and egg abnormality, we used linear mixed effects models (*lme* in *lme4*: Bates et al. 2015) with bleaching history and year as fixed effects, and colony ID as a random effect. Analysis of variance (ANOVA) tables were generated using type II sum of squares (*Anova* in *car*: Fox and Weisberg 2011).

To test the effects of parental bleaching history on offspring performance, we first analyzed the proportion of eggs fertilized and the proportion of eggs reaching each developmental stage 3-hours post fertilization using linear mixed effects models with cross-type as a fixed effect and colony ID of egg donor and sperm donor as random effects. We then analyzed the proportion of larvae that settled at 7, 28, and 59-days post-fertilization using a linear mixed effects model with cross-type and day (7, 28, and 59-d post-fertilization) as fixed effects and colony ID of egg donor and sperm donor as random effects. Lastly, we generated survivorship estimate curves to visualize offspring fate by cross-type with *ggsurvplot* of the census over time (i.e., days 6, 7, 23, 27, 28, 53, and 59 post-fertilization) (*survfit* in *survminer*; Kassambara et al., 2017). Cox proportional hazards (CPH) model was used to analyze the effects of cross, egg donor, and sperm donor individually on offspring survivorship (*coxph* in *survminer*; Kassambara et al., 2017). Dispersion parameters were inspected through a simulation-based approach (*DHARMa* package: Hartig, 2019).

## RESULTS

### 3.1 Sexual reproduction and egg traits

All fifteen colonies observed in this study released egg-sperm bundles one or more nights in both years (Figure 3A). When spawning was observed, colonies began releasing egg-sperm bundles between 20:20 and 21:32 hrs and ended between 20:30 and 22:15 hrs. Parental bleaching history did not affect the occurrence of spawning (*P* = 0.619) and had no interactive effect with year (*P* = 0.982). The proportion of colonies releasing gametes significantly differs by year (*P* < 0.001). In 2017, the proportion of colonies participating in spawning events was 36% lower than in 2016. In both years, the second month of the spawning season had the highest proportion of colonies spawning.

In 2016, the spawning season following consecutive bleaching events, colonies that bleached and recovered had 22.5% higher mean total reproductive output than colonies that did not bleach, although this was not statistically significant (Fig. 3B; Table 1). There was no effect of year and no interaction between bleaching history and year or reproductive output (Figure 3B; Table 1).

Individual egg volume ranged from 0.032 to 0.099 mm^3^ and did not differ by parental bleaching history, year, or by their interaction (Figure 3C; Table 1). The number of eggs per bundle from both bleached and nonbleached parental colonies ranged from 2 to 29, and mean eggs per bundle for all colonies examined was 13.3% less in 2016 than in 2017 (Figure 3D; Table 1). Eggs per bundle did not differ by parental bleaching history (Table 3). There were 79.5% more eggs with irregularities in 2016 than in 2017 with no difference by bleaching history (Figure 3E; Table 1).

### 3.2 Fertilization, survivorship, and settlement

While reproduction continued in the colonies examined, we found that cross-type did influence the fertilization, embryonic development, and percent motile larvae (Figure 4A; Table 2).

**Figure 4.**
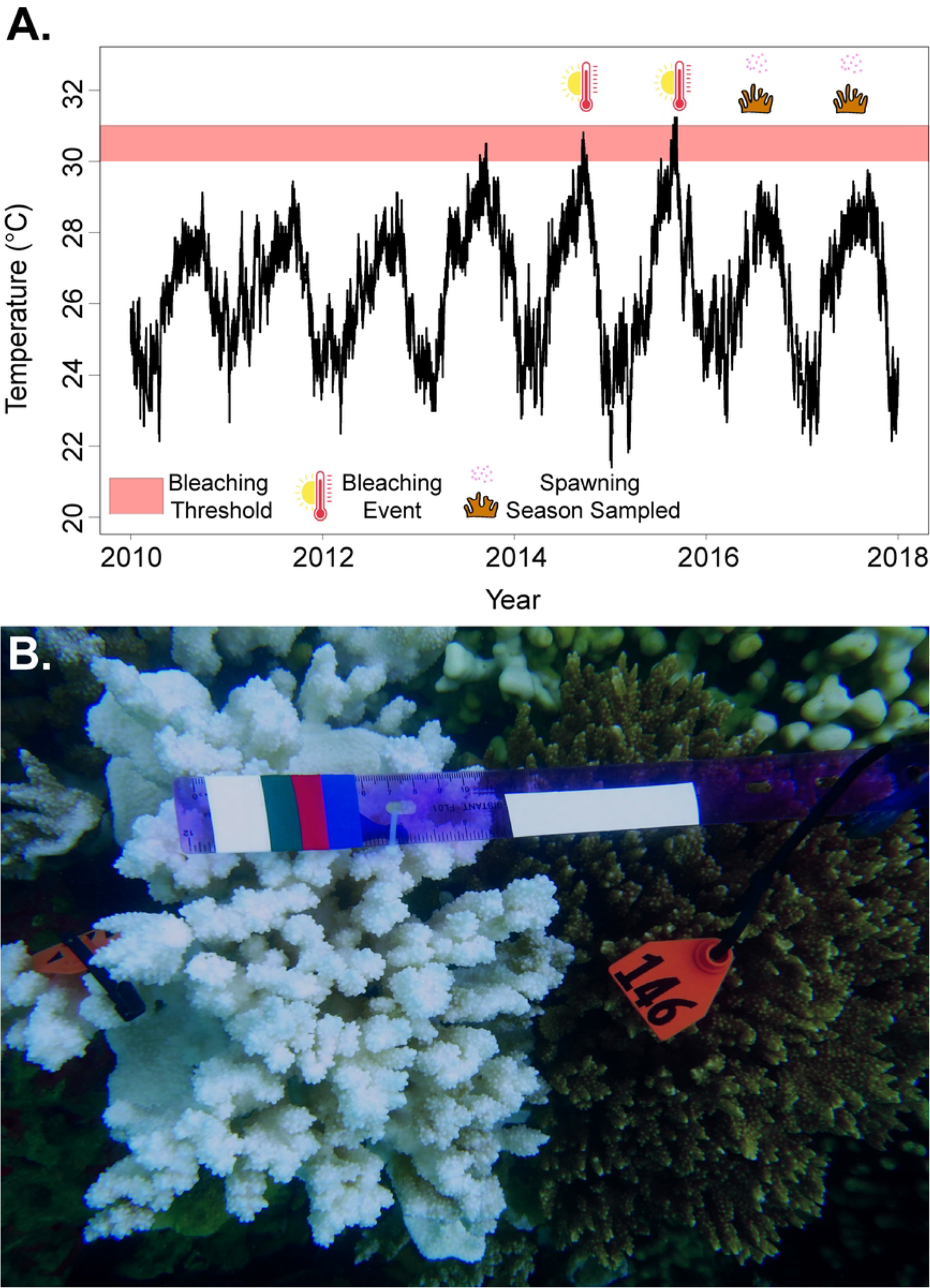
Offspring performance from selected crosses. A) Images of fertilized eggs and embryos (scale bar = 500 μm), planula larvae (scale bar = 500 μm), and settlement (1 mm). Mean ± SE. B) proportion of eggs fertilized by cross-type, C) proportion of cell division after 3-h fertilization D) proportion of swimming larvae and E) settlers during five timepoints over a 59-d period, and F) survivorship estimate curves by cross over seven timepoints between 6 and 59-d with the figure embedded comparing the survivorship curves of offspring from bleached and nonbleached egg donors.

Fertilization success differed by cross-type (Figure 4B; *P* = 0.033), with higher fertilization success in the NB × NB cross-type than in the B × NB and NB × B cross-types although this was not statistically significant (*P* < 0.080). There were no differences between B × B and the other three cross-types (*P* ≥ 0.342). Cell division advanced more quickly for within-type crosses (B × B and NB × NB) than between-type crosses (B × NB and NB × B) at 3-h post-fertilization. Embryos from B × B and NB × NB reached the 16-cell stage at 3-h post fertilization, whereas embryos from B×NB and NB × B crosses were only at the 4-cell stage (Figure 4C, Table 2).

Percent swimming larvae and settlement varied by cross-type, driven by egg donor bleaching history (Figure 4D/E; Table 3). Offspring developed from eggs from previously B egg donors had lower survivorship than those from NB egg donors. NB egg donors had a significant effect on the proportion of swimming planula larvae (Figure 4E; *P* < 0.001). However, no difference was found in offspring survivorship from bleached or nonbleached sperm donors (*P* = 0.992). Overall, percent mortality from the initial to final time point (i.e., day 5 to 59) were 92.5% for B × B, 87.8% for B × NB, 85.6% for NB × B, and 77.3% for NB × NB (Figure 4F).

## DISCUSSION

Unprecedented back-to-back warming events (2014 and 2015) in Kāneʻohe Bay, Hawaiʻi influenced the reproductive capacity of *M. capitata*. In the first spawning season after the 2015 bleaching event, *M. capitata* had eggs with similar volume but fewer eggs were packaged within bundles. In 2016 egg abnormality was higher than in 2017 regardless of parental bleaching history. These results demonstrate the impacts of marine heatwaves and coral bleaching on reproductive capacity in the following spawning season with the capacity to recover in subsequent nonbleaching years. Delayed beneficial maternal effects observed in larvae were driven by corals resistant to bleaching. These results demonstrate that although *M. capitata* has the energetic capacity to continue reproduction despite bleaching response, cross-generational impacts occur (Byrne et al., 2020), with possible ecological consequences.

### 4.1 Reproductive capacity after bleaching events

*M. capitata* appears to maintain reproductive resilience after consecutive marine heatwaves and coral bleaching events, as evidenced by continuing synchronous broadcast spawning and production of viable eggs and sperm. These results are consistent with prior studies examining the influence of environmental and biological factors on *M. capitata* gametogenesis and spawning in Kāneʻohe Bay (Cox, 2007; Padilla-Gamiño et al., 2014). For instance, Padilla-Gamiño et al. (2014) found similar rates of gametogenesis along a strong sedimentation gradient. Further, Cox (2007) found no differences in reproductive output, eggs per bundle, and egg size between B and NB parents in the spawning season immediately following the 2004 mild warming event. Resilience in *M. capitata* may be due to its capacity to maintain energetic stability under stress (Wall et al., 2019), here evident by the completion of gametogenesis even at the cost of producing fewer eggs per bundle with higher proportion of irregularity in shape in 2016 than in 2017. One hypothesis to explain similar reproductive traits in bleached and nonbleached parents, is that after the thermal stress (Sept-Oct), there is still time for the colonies to recover (∼5-6 months) and develop gametes that can be released during the spawning season (May-August).

Maintaining egg traits such as size and biochemical composition would serve as an advantageous strategy to ensure ecological fitness of parents and their developing offspring (Moran & McAlister, 2009; Padilla-Gamiño et al., 2013; Jacobs & Podolsky, 2010). For example, there may be an optimal egg size that needs to be achieved to ensure successful fertilization (Levitan, 1993; Levitan & Petersen, 1995). It is notable that the relationship between egg size and number of eggs per bundle in our study has shifted from prior studies; we found 10-12 eggs per bundle in 2016-2017 compared to 15-18 eggs per bundle in studies prior to 2010. (Cox, 2007; Padilla-Gamiño & Gates, 2012), and egg size was 11% larger in our study than previous studies. This apparent tradeoff in reproductive effort suggests plasticity in response to environmental changes and emphasizes the need for long-term studies to detect changes in sexual reproduction (Levitan et al., 2014; Hagedorn et al., 2016; Price et al., 2019; Schlesinger et al., 2019).

High inter- and intraspecific variation in thermal tolerance contribute to reproductive consequences after bleaching events (Baird & Marshall, 2002; Levitan et al., 2014; Fisch et al., 2019). For example, there were no differences in percent reproductive polyps between bleached and nonbleached colonies of acroporid species at Heron Island on the Great Barrier Reef after the 1998 bleaching event (Ward et al., 2000). Baird and Marshall (2000) found that the bleaching response of *Acropora millepora* did not influence fecundity, whereas the bleaching response of *Acropora hyacinthus* strongly influenced the completion of gametogenesis. It is important to emphasize that although reproductive capacity after bleaching events can be greatly suppressed, there are species and populations that are resistant and/or more able to recover from bleaching (Szmant & Gassman, 1990; Omori et al., 2001; Baird & Marshall, 2002; Cox, 2007; Levitan et al., 2014; Ward et al., 2000). Exceptional populations carrying resilient individuals are critical to identify and protect, particularly if they are successful in continuing sexual reproduction to replenish impacted neighboring reefs (Underwood et al., 2007; Baker et al., 2008). Coral reproductive modes and strategies have evolved to withstand environmental fluctuations and severe selective pressures, but the question of how much thinning can a population withstand without complete collapse remains.

### 4.2 Parental effects on fertilization and offspring survivorship

We demonstrate maternal effects, or cross-generational plasticity, due to bleaching resistance in *M. capitata*, with effects apparent at fertilization and through offspring survivorship. Fertilization success differed by cross-type which may be due to gametic compatibility (Vacquier, 1998). Such compatibility could be driven by gamete-recognition proteins that mediate fertilization through chemoattraction, binding, and fusion of egg and sperm (Vacquier, 1998; Tomaiuolo & Levitan, 2010; Miller et al., 2018). Furthermore, high gamete compatibility may explain the advanced rate in cell division during embryogenesis in offspring from NB × NB and B × B cross-types. Egg-sperm compatibility has been observed as a mechanism for pre-zygotic isolation to select for populations that are likely to succeed under intense environmental pressures, such as temperature (Coll et al., 1994; Baums et al., 2013; Kosman & Levitan, 2014; Vermeij & Grosberg, 2018). With regards to sperm selection, Henley et al. (2021) demonstrated sperm motility in *M. capitata* is strained with a severe decline that may be associated with damaged mitochondria in response to heat stress. In this study, bleached colonies had similar fertilization success to nonbleached colonies, the thermally tolerant individuals will likely have a stronger selective advantage as warming continues to weed out thermally sensitive adults and their offspring (Drury et al., 2022).

Eggs from parent colonies that were resistant to bleaching had offspring with notably higher survivorship regardless of the sperm donor bleaching history. More pronounced benefits of nonbleached egg donors support previous work of maternal provisioning in coral offspring (Marshall et al., 2008; Quigley et al., 2016; Chan et al., 2019). Beneficial cross-generational plasticity through maternal effects observed in offspring survivorship may be attributed to energetic provisioning through lipid reserves stored in the eggs and larvae (Jones et al., 2011; Padilla-Gamiño et al., 2013; Rivest et al., 2017), mitochondria (Dixon et al., 2015), or vertical transmission of Symbiodinaceae from the parent into the eggs (Jones et al., 2010; Padilla-Gamiño et al., 2012; Quigley et al., 2016).

*M. capitata* houses the endosymbionts *Cladocopium* spp. and *Durusdinium* spp., formerly Clade C and D, respectively. It has been shown that *M. capitata* colonies associate with *Durusdinium* spp. in more challenging environments such as high light and variable thermal regimes (Padilla-Gamiño et al., 2013, but see Stat et al. 2011). During heat wave events, colonies that were resistant to bleaching were associated with a host mixture of *Cladocopium* spp. and *Durusdinium* spp. while susceptible colonies to bleaching were dominated by *Cladocopium* spp. (Cunning et al., 2015). Colonies resistant to bleaching have also shown distinct metabolomic signatures mainly driven by betaine lipids from the symbionts (Roach et al. 2021). In *M. capitata*, these symbionts are vertically transferred to the eggs creating offspring with different assemblages (Padilla-Gamiño et al., 2013) that could confer different physiological attributes to the offspring. For example, Little et al. (2004) found that *Acropora* juveniles grew faster when infected with clade C than clade D, and Abrego et al. (2008) showed enhanced physiological tolerance and higher ^14^C photosynthate incorporation in juveniles infected with clade C1. Padilla-Gamiño et al. (2013) showed that *Cladocopium* spp. is more likely to be transferred to *M. capitata* eggs, but further research is needed to better understand transfer mechanisms, and how different symbionts influence survival, tolerance and/or tradeoffs in larvae and juveniles. Our results suggest that nonbleached colonies have higher ecological fitness than their bleached counterparts and may benefit the maintenance of genetic diversity while guiding populations towards higher thermal tolerance in a warming ocean.

### 4.3 Interventions for thermal tolerance

Research on coral reefs has become greatly focused on identifying human interventions (i.e., assisted evolution) that support biological persistence and resilience against anthropogenic stressors (van Oppen et al., 2015, 2017; National Academies of Sciences et al., 2019).

Developing effective interventions to implement has become increasingly urgent to protect shallow-dwelling coral reef ecosystems (NASEM, 2019). Current strategies proposed to overcome bottlenecks in early life history include identifying genetic adaptation (Dixon et al. 2015), environmental hardening through non-genetic or epigenetic mechanisms (Putnam & Gates 2015; Putnam et al. 2020, 2016; Liew et al. 2020; Dixon et al. 2018), manipulation of coral-algal endosymbiosis (Chakravarti & van Oppen 2018; Levin et al., 2017; Buerger et al., 2020; Dilworth et al., 2021), cryopreservation for coral conservation (Hagedorn et al., 2017), and selective breeding (Quigley et al., 2016; Chan et al., 2018, 2019).

Human interventions applying selective breeding in coral sexual propagation has been proposed as one of the viable options to maintain genetic diversity and increase resilience in restoration efforts (Barott et al., 2021; Epstein et al., 2003; van Oppen et al., 2015, 2017; NASEM, 2019; Hancock et al., 2021); however, feasibility to potentially scale up efforts remain limited and costly without full understanding of tradeoffs (Edwards et al., 2015; Chamberland et al., 2017). Our study supports the potential for selective breeding and environmental hardening to have positive fitness consequences. In our study, bleaching in *M. capitata* did not disrupt reproductive output or physical traits of the eggs (size and abnormality), but the use of eggs from NB colonies in the intentional crossing of gametes produced offspring with higher settlement and survivorship, while bleached corals had higher overall fecundity to balance reduced survivorship and settlement. These results are important to maximize restoration efforts through selective breeding by identifying candidate colonies in the natural environment or through manipulated stress tests. We encourage further research to test the efficacy and trade-offs of human-assisted evolution, particularly selective propagation and environmental hardening, designed to increase coral resistance that would ensure the continuation of coral reefs confronted by global climate change.

## ACKNOWLEDGEMENTS

This work was supported by funding from Paul G. Allen Family Foundation to RG and the National Science Foundation Graduate Research Fellowships to EAL. We would like to thank members of the Gates Coral Laboratory for their technical support and advice, especially Jen Davidson and Dr. James Guest, we are grateful to Mary Hagedorn, Amy Moran, and Peter Marko for their feedback on the manuscript, the many volunteers especially Dyson Chee, Megan Buras, Katie Allen, Shayne Fabian, and Kat McPherson and the security Greg Miranda and Moses at Moku o Loʻe who ensured safety and the success of this research. We dedicate this research to Dr. Ruth D. Gates and her infectious enthusiasm that pushed human-assisted evolution to the forefront of coral biology – you rock!

## TABLES

**Table 1.** Statistical summary of Type II Wald χ^2^ test of generalized linear mixed effects model and linear mixed effect models testing the fixed effects of spawning year and parent history of bleaching susceptibility on sexual reproduction and offspring performance. Significance indicated in bold text.

**Table 2** Statistical summary of Type II Wald χ^2^ test of linear mixed effects model testing the fixed effects of cross-type on the proportion of fertilized embryos, cellular development, and larvae and settlers over three-timepoints. Significance indicated in bold text.

**Table 3** Summary of Cox proportional hazards (CPH) analysis of coral offspring survival influenced by the fixed effects: cross-type, egg donor, and sperm donor over time with model average estimates of the hazard ratio (with 95% confidence intervals; Cross: df = 3 or Egg/Sperm donor: df = 1; *n* = 1,318; number of events = 560). Bold text indicates significance.

